# The archaeological potential of the northern Luangwa Valley, Zambia: the Luwumbu basin

**DOI:** 10.1101/2022.05.18.492447

**Authors:** A. Burke, M. Bisson, F. Schilt, S. Tolan, M. Drapeau, J. Aleman, M. Peros

## Abstract

The Luangwa Basin, Zambia, which forms part of the Zambezi drainage, is strategically located between the Central African plateau and the East African Rift system. The Luangwa River and major tributaries, such as the Luwumbu River, are perennial water sources supporting essential resources that sustain human communities and a rich and diverse fauna and flora. The archaeological record of Luangwa is relatively unknown, despite early archaeological exploration hinting at its potential. Recent research in the southern Luangwa valley, however, suggests that it preserves a long record of hominin occupation spanning the Early to Late Stone Age. The research described here details fieldwork carried out in northeastern Luangwa, in the Luwumbu Basin, that confirms that a relatively deep package of Quaternary deposits, containing evidence of the Stone Age occupation of the region persists in the upper piedmont zone.

## Introduction

The Luangwa Valley is a lateral extension of the East African Rift system, cutting diagonally through the high central plateau of Zambia (1). The valley is known for its rich paleontological deposits, which have been instrumental in refining our understanding of the impact of the end Permian extinction and the subsequent evolution of vertebrate communities during the Triassic (2, 3). Despite its proximity to the East African rift, with its long history of archaeological investigation, the archaeological record of the Luangwa Valley is still largely unexplored.

Early reports of Stone Age material in surface deposits in the Luangwa Valley date to the first half of the 20th Century (4, 5). Archaeological research in the region was not pursued further at the time, due to a lack of chronostratigraphic information and access difficulties (6, 7). In the late 20^th^ Century hydrocarbon exploration, advances in satellite imagery and the development of wetland management schemes contributed to renewed scientific interest in the hydrology and sedimentary history of the Luangwa Basin, e.g., (8–11). New archaeological and geoarchaeological explorations of the southern Luangwa valley began in 2003, confirming the presence of Stone Age and Iron age material, occasionally in stratified contexts (6, 12, 13).

Here, we report on archaeological fieldwork conducted in northeastern Luangwa in 2016 and 2019, more precisely in the piedmont zone within the Luwumbu sub-basin and part of the Viziba sub-basin (Fig 1). A more limited exploratory survey conducted in the central part of the Luangwa basin is not reported on here. The results of this research are comparable with research conducted in central and southern Luangwa (12, 13) and in the Karonga district, northern Malawi (14), on the other side of the divide between the Luangwa Valley and Lake Malawi, allowing us to draw a more complete picture of the archaeological potential of the Luangwa Basin.

**Fig 1.**
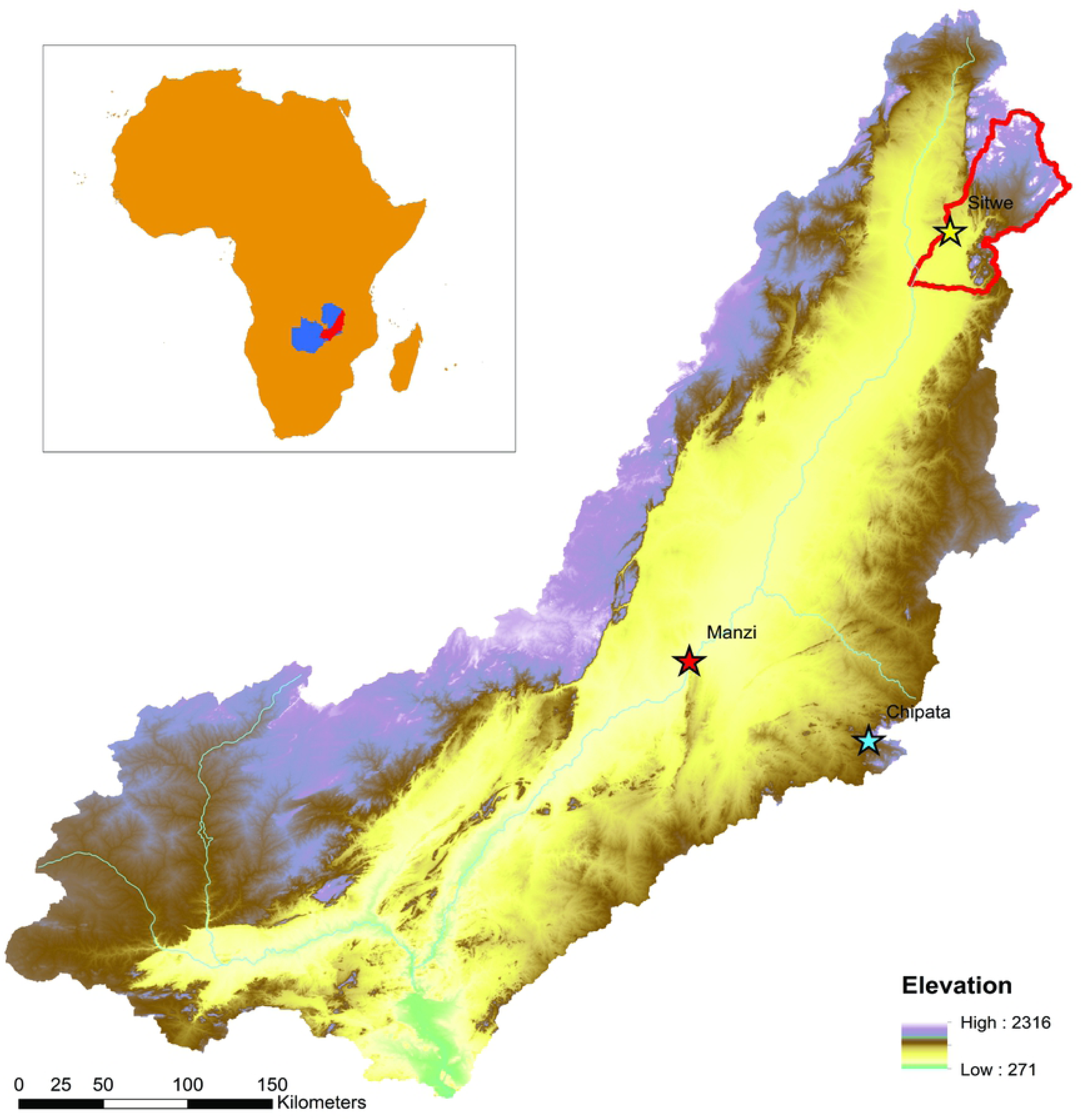
The Luangwa Valley, Zambia. The fieldwork location in the piedmont zone of northeastern Luangwa is highlighted in red.

### Geography of the Luangwa Basin

The Luangwa Basin, northeastern Zambia, borders Malawi, Mozambique and Zimbabwe. The basin has an area of 145,690 km^2^ and is a sub-catchment of the Zambesi, a major hydrological system that drains a large portion of south-central Africa before discharging into the Indian Ocean off the coast of Mozambique (15, 16). The valley is drained by the Luangwa River, which flows 850 km in a southwesterly direction from its headwaters in the Mafinga Hills to its’ confluence with the Zambezi. As a perennial water course, the Luangwa River attracts diverse and abundant wildlife, especially during the dry season, and sustains local farming communities in addition to supporting hunting and tourist industries.

The Luangwa Basin ranges from 2,297 meters asl at its’ headwaters to 323 meters asl in the southern floodplain (Fig 1). Average annual precipitation in the valley, calculated over a 20-yr period, is 966 mm/year (Fig 2a), mostly falling during the rainy season, from December to March. During the dry season rainfall virtually ceases, although negligible amounts of rain continue to fall in the wetter, northern sub-basin. Regional vegetation cover is dominated by grassland and deciduous miombo woodland (Fig 2b). The dynamic floodplain widens in the central portion of the basin where the Luangwa River’s main channel forms a network of meanders that actively reshape the landscape (11).

**Figure 2:**
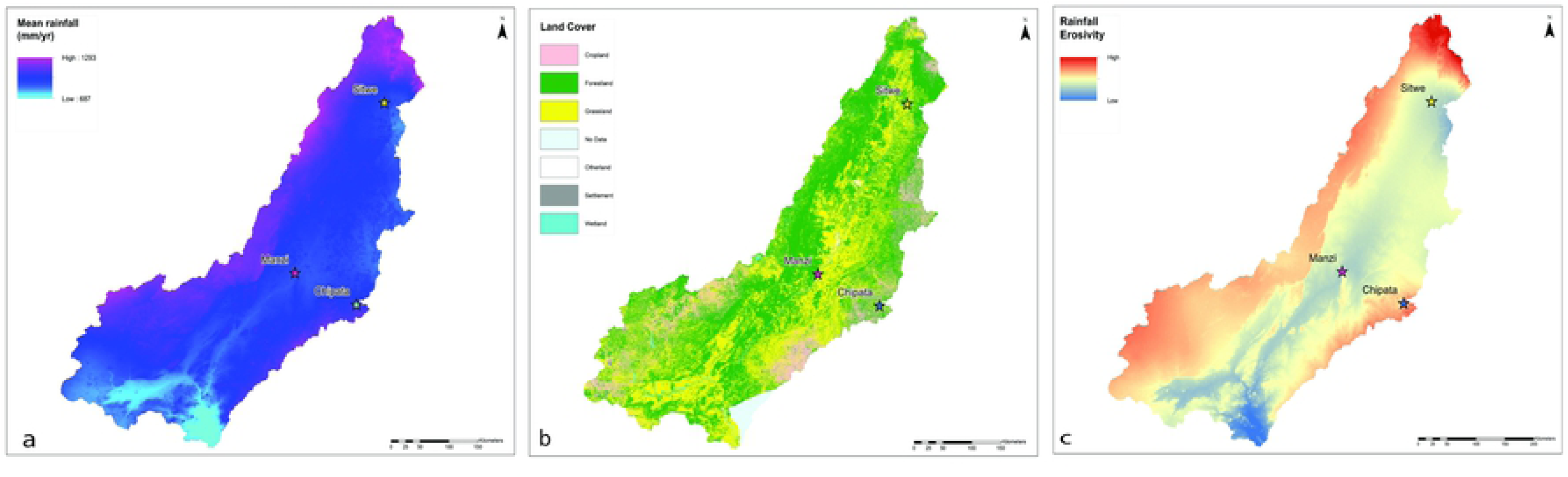
Geography of the Luangwa Basin. Clockwise from top left: a) rainfall patterns; b) landcover; c) rainfall erosivity.

#### Structural setting

The Luangwa Valley is a Karoo rift basin, located on the southeastern border of the Central African Plain, an extensive erosion surface comprised of residual soils and laterite that forms a high plateau at the heart of the continent roughly 1200 m asl (1). The basin forms a distinct geosystem (9) with well-defined structural units: two opposing half-grabens define a southern and a northern sub-basin, separated by a central transfer zone just south of 12°S (10). The major boundary fault of the northern half-graben is located on the eastern border of the basin and the situation is reversed in the southern sub-basin. The Luangwa basin was further shaped by tectonic activity along the Mwembeshi Shear Zone, which traverses the southern sub-basin from east to west, during the early Permian to early Triassic with the strongest impact in the southern sub-basin (10).

#### Geological setting

The geological formations of Eastern Zambia were initially described in the first half of the last century (17) with a focus on Paleogene/Neogene formation processes. Subsequent fieldwork in the Luangwa Basin focussed on the early sedimentary sequence and the rich, Permo-Triassic, fossil-bearing deposits, e.g., (18). Geological surveys conducted at the end of the 20^th^ century to assess the hydrocarbon potential of the region employed extensive seismic studies, gravity and aeromagnetic surveys, and deep exploration wells to define the sedimentary sequence, producing the structural model presented above (10). Geomorphological fieldwork in the eastern foothills, near Chipata (8, 9) (Fig 1), paleontological fieldwork in the northeast, e.g., (2, 19, 20), and geoarchaeological fieldwork in the southern sub-basin, in the vicinity of Manzi (12, 13, 21) (Fig 1) further our understanding of the Luangwa Valley’s sedimentary history.

The sedimentary sequence in the Luangwa Basin consists of a succession of Permo-Triassic deposits (the Lower and Upper Karoo) up to 8000 m thick in the centre of the basin. These deposits are overlain unconformably by the Luangwa formation, a post-Karoo formation consisting of poorly consolidated sandstones, lacking biostratigraphic controls, which could date to the Cretaceous or the Paleogene (10). The “post-Karoo” sediment burden is considerably lighter in the northern sub-basin. Overlying the Luangwa formation is a succession of poorly differentiated Neogene deposits capped by Quaternary sediments, composed of colluvium and alluvium, which have been substantially reworked in places by climate-driven changes to the hydrological system (11).

#### Archaeological record of the Luangwa Basin

Surface discoveries of Stone Age artefacts in the Luangwa Valley have been reported since the early 20^th^ century (4, 5). Archaeological fieldwork conducted subsequently in southern Luangwa, extending as far as Kamnama, ca. 25km north of Chipata near the Zambia/Malawi border, allowed Phillipson to suggest a regional sequence of Stone Age industries (22) which begins with the Middle Stone Age (MSA) (mode 3), followed by a succession of microlithic Late Stone Age (LSA) industries and finally, a succession of Iron Age industries (Op. Cit.). More recently, on the southern floodplain of the Luangwa River, in the southern sub-basin, Early Stone Age (ESA) (Mode 1) cores were discovered on a sandbar in conjunction with fossil vertebrates (23). In the foothills of northern Luangwa, Macrae and Lancaster (1937) reported a chronologically mixed assemblage of stone tools in exposed gravel deposits on the banks of the Viziba, a tributary of the Luangwa River. In 1963, geologists working in the piedmont of northeastern Luangwa, confirmed the presence of MSA (Mode 3) tools in superficial contexts (18).

In 2003, a 5-year programme of archaeological survey and exploration began focussing on the transfer zone between the Nchindeni Hills (which rise up in the centre of the transfer zone) and the Muchinga escarpment, on the western edge of the Luangwa Basin. This research confirmed the presence of ESA, MSA and LSA artefacts in superficial deposits and in stratified sedimentary contexts, significantly contributing to our understanding of the archaeology of the Luangwa Valley while highlighting the difficulty inherent in obtaining direct dates from contexts in the dynamic floodplain environment (6, 12, 13, 24). Our fieldwork seeks to complete the picture provided by existing research in southern Luangwa, adding to our appreciation of the archaeological potential of the region.

### The study region: Northeastern Luangwa Valley

#### Geographical setting

We conducted archaeological fieldwork in the foothills of northeastern Luangwa in 2016 and 2019, concentrating on a heavily eroded and dissected piedmont zone, culminating at ca. 900 m asl, that forms the divide between the northern Luangwa and Luwumbu sub-basins (Fig 3a). The piedmont surface consists of a series of sandstone outcrops separated by superficial deposits of sandy and lateritic sediment with pockets of fossil wood. Quartzite cobbles are ubiquitous, occurring on exposed hilltops, slope deposits and in river-bed deposits. Cobble levels of variable thickness are visible in sediment exposures and stream-cut banks. Cobbles consist predominantly of quartzite and white quartz, with occasional blocks of fossil wood and, much more rarely, small and irregular flint nodules.

**Figure 3:**
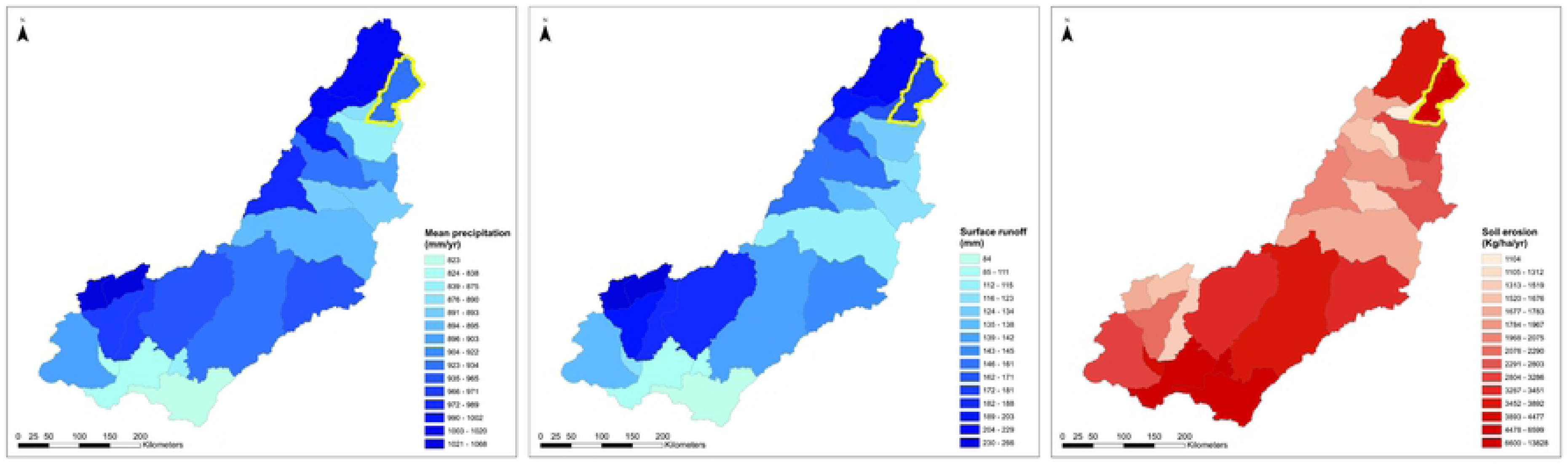
Geography of the study region: the piedmont zone. Annual precipitation (a) and surface runoff (b) for sub-basins. The Luwumbu catchment is highlighted in light blue.

The Luwumbu catchment, which is the main focus of this research, has an area of 4476 km^2^ and is fed by multiple streams and rivers. The Luwumbu is a perennial river that flows from the Mafinga Hills, where the headwaters of the Luangwa River also form. The Luwumbu sub-basin drains the Makutu Range, a high relief mountain chain (elev. ∼1700 m asl) that extends southward from the Mafingi Hills, to the west, and the Nyika Plateau to the east. The Luwumbu is joined by the Lusangani River at the foot of the Ngonya Hills near Sitwe village, where it forms a broad floodplain before flowing southwest to its’ confluence with the Luangwa River ∼45 km downstream. A series of small agricultural communities, including Sitwe, are situated in the floodplain and along the Luwumbu and the Lusangani. Land-use patterns in the region affect the distribution of vegetation cover, which in turn influences patterns of soil erosion and shapes the hydrological system (25, 26). Vegetation cover is dominated by miombo woodland in the foothills, with a mosaic of fallow land, cropland, and grasslands in the river valleys (Fig 3).

High seasonal rainfall in the region (Fig 3) contributes to active soil erosion, localised gullying and incision of the piedmont surface. Annual soil loss estimates for the Luwumbu catchment (and the adjacent Viziba sub-catchment, to the west) is on the order of 12,000 – 22,000 tons/km^2^/annum (USDA 2011) and erosivity estimates for the mid-course of the Luwumbu catchment, where fieldwork took place, are high due to a combination of topography, soil and cover types, and rainfall patterns (15).

#### Geological setting

The piedmont zone in northeastern Luangwa forms part of the Upper Grit Formation, which consists of red-brown pebbly feldspathic sandstone with trough and planar cross bedding, further described as containing fossil wood (27). The Upper Grit unconformably overlies the Red Marl Formation which outcrops ca. 3 km north of the main survey area. The Red Marl Formation is an Upper Karoo formation consisting of siltstone and fine-grained arkosic sandstone (at least 25% feldspar). Other Upper Karoo Formations in the Luwumbu Basin include the fossil-rich siltstones and mudstones of the Ntawere Formation (S2), and sandstone and feldspar-rich sandstone (arkose) of the Escarpment Grit Formation (S1). Older Lower Karoo siltstones and mudstone of the Madumabisa Mudstone Formation (R3 and R3’) are exposed in the north and east of the study region. Quaternary deposits are not mapped.

The Makutu Range, north of the piedmont, forms the western margin of the Luwumbu sub-basin. Formed of metasedimentary rock correlated with the Mafingi Group, the lithology of the Makutu Range consists mainly of sericite schists, quartz phyllites, white fine grained orthoquartzite bands, and quartzites overlying basal granite (27–29). Superficial deposits of what O’Connor described as “sandy to lateritic colluvium” are observable in the northern part of the catchment, where he also describes “blankets of Makutu quartzite boulders” (O’Connor 1999:8) on the remains of an erosion surface hypothetically formed during the Tertiary (28).

#### Archaeological record

Previous surveys in the foothills of northeastern Luangwa record the presence of fossil wood and fossil vertebrates in numerous locations in the piedmont zone separating the Luangwa and Luwumbu sub-basins, in Lower and Upper Grit exposures (30). Subsequent paleontological fieldwork conducted in the Luwumbu catchment for the Field Museum, Chicago, from 2009 to 2014, confirmed the presence of rich fossil deposits in a series of exposures of the Ntawere formation (Upper Grit), e.g., (3, 19, 20, 31, 32). Reconnaissance surveys undertaken by ST also revealed the presence of concentrations of Stone Age artefacts at several localities on the margins of the upper piedmont surface, within the Luwumbu catchment. These localities form the focus of the archaeological fieldwork we conducted in the region in 2016 and 2019 with the aim of establishing its’ archaeological potential [FIG 4].

**Fig 4.**
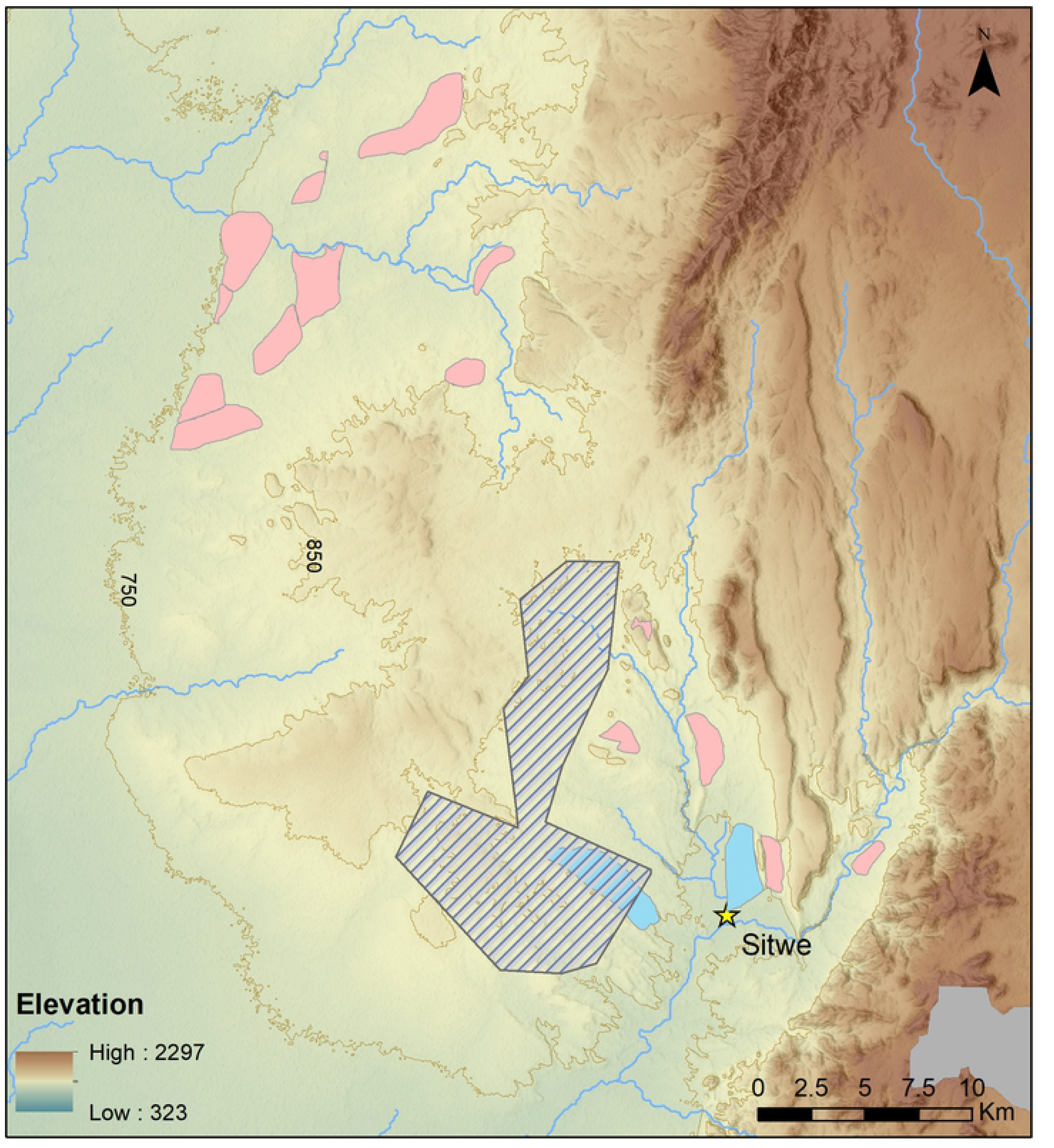
Survey zones. Drysdale & Kitching (1963) survey zones courtesy of Neil Tabor (Pink: Triassic, Blue: Upper Permian). Cross-hatched: Phase 1 Survey zones (this project).

## Methods

### Archaeological surveys

Pedestrian surveys were conducted in two phases: exploratory surveys (Phase 1) and archaeological surveys (Phase 2). Limitations encountered included difficulty of access to the survey region, which lacks a road system, and the absence of power sources, cellular coverage or internet access.

©Garmin CX GPS devices were used to record locations and artefact positions using a “catch and release” strategy in Phase 1 (exploratory surveys) and Phase 2 surveys carried out in 2016, when the techno-typological identification of artefacts was recorded in the field. Phase 2 surveys conducted in 2019 adopted a “tag and bag” strategy and lithic artefacts were transported to the University of Lusaka for metrical, techno-typological and taphonomic analysis. GPS data was downloaded daily and processed using ©Garmin Basecamp software. Surveys took place in August at the height of the dry season, when surface visibility is good to fair in the upland region due to the withering of water-stressed vegetation. Clear skies and sparse tree cover meant the accuracy of the hand-held GPS devices averaged ∼2-3 m, not exceeding 5m.

Phase 1 (exploratory) surveys focus on the Luwumbu catchment, encompassing the piedmont zone and adjacent, upstream locations associated with the Upper and Lower Karoo formations (Fig.5). Surveys were conducted by an avocational archaeologist (ST) in 2014 and 2016 targeting sediment exposures identified and numbered prior to fieldwork using satellite imagery (Google Earth). The spatial coordinates of the locations explored during Phase 1 were recorded and the presence/absence of artefacts, fossil wood and/or fossil vertebrates was noted. Additional Phase 1 exploratory surveys were conducted in 2019 by a small, interdisciplinary team (3-4 people) on the margins of the upper piedmont, from the headwaters of the Lutete and the Nkande rivers to the headwaters of the Viziba River, and in the valley floors along the upper reaches of the Lutete and the Nkande. Typologically distinct tools encountered during these surveys were georeferenced and photographed. A total of 63 locations were investigated during Phase 1, lithic samples from two locations were described in the field (MO1, SW22) and five locations were identified as localities of interest due to the presence of relatively dense lithic scatters and earmarked for Phase 2 (SURVEY 1, SW23, SW23A, SW60 and SW37) (Fig.6).

**Fig 5.**
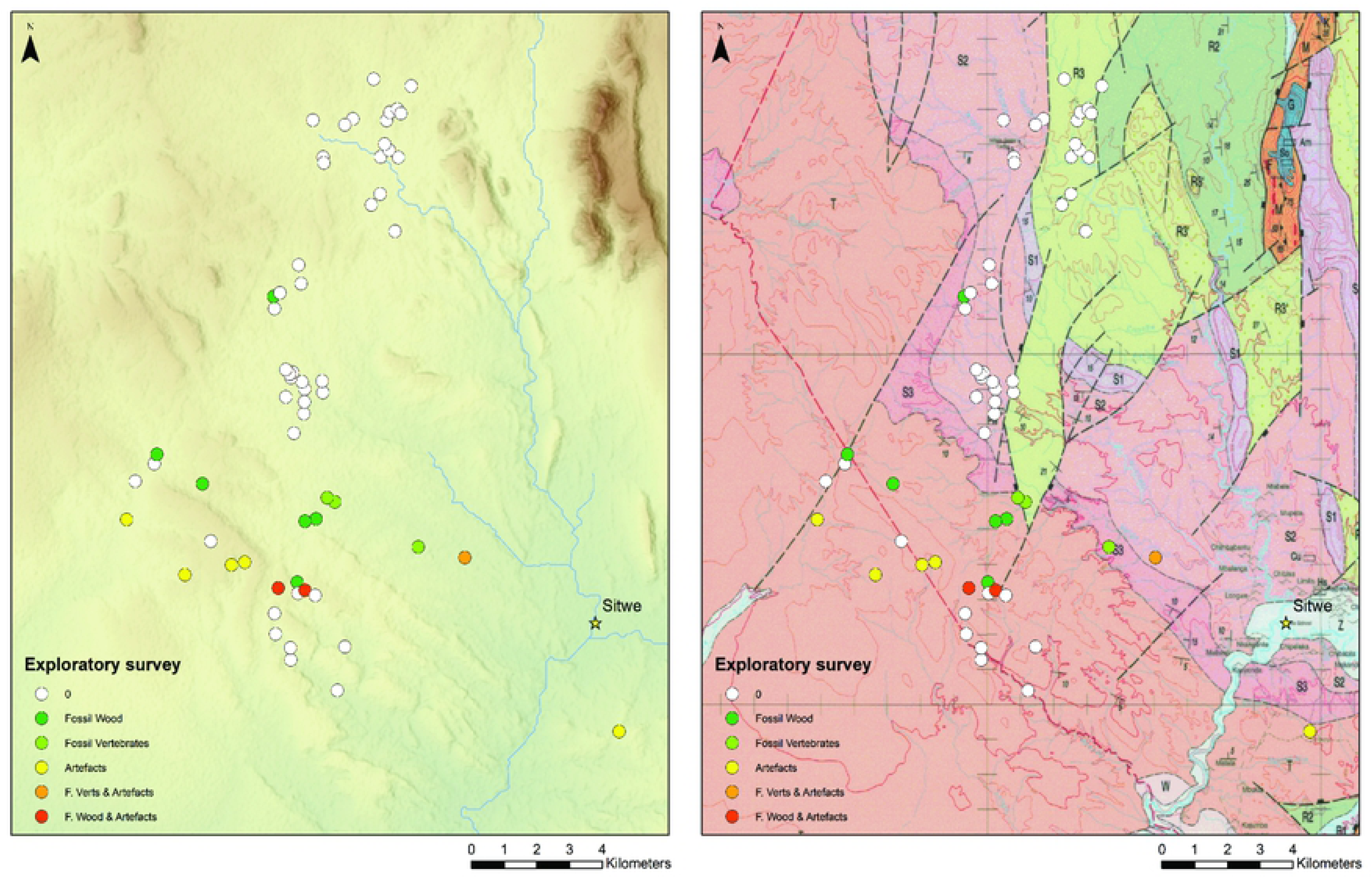
Phase 1 exploratory survey locations. The nature of finds at each location is colour coded (see legend). Geological map (right): Upper Grit Formation (T), Ntawere Formation (S3, S2), Madubisa Mudstone formations (R3, R2).

**Fig 6.**
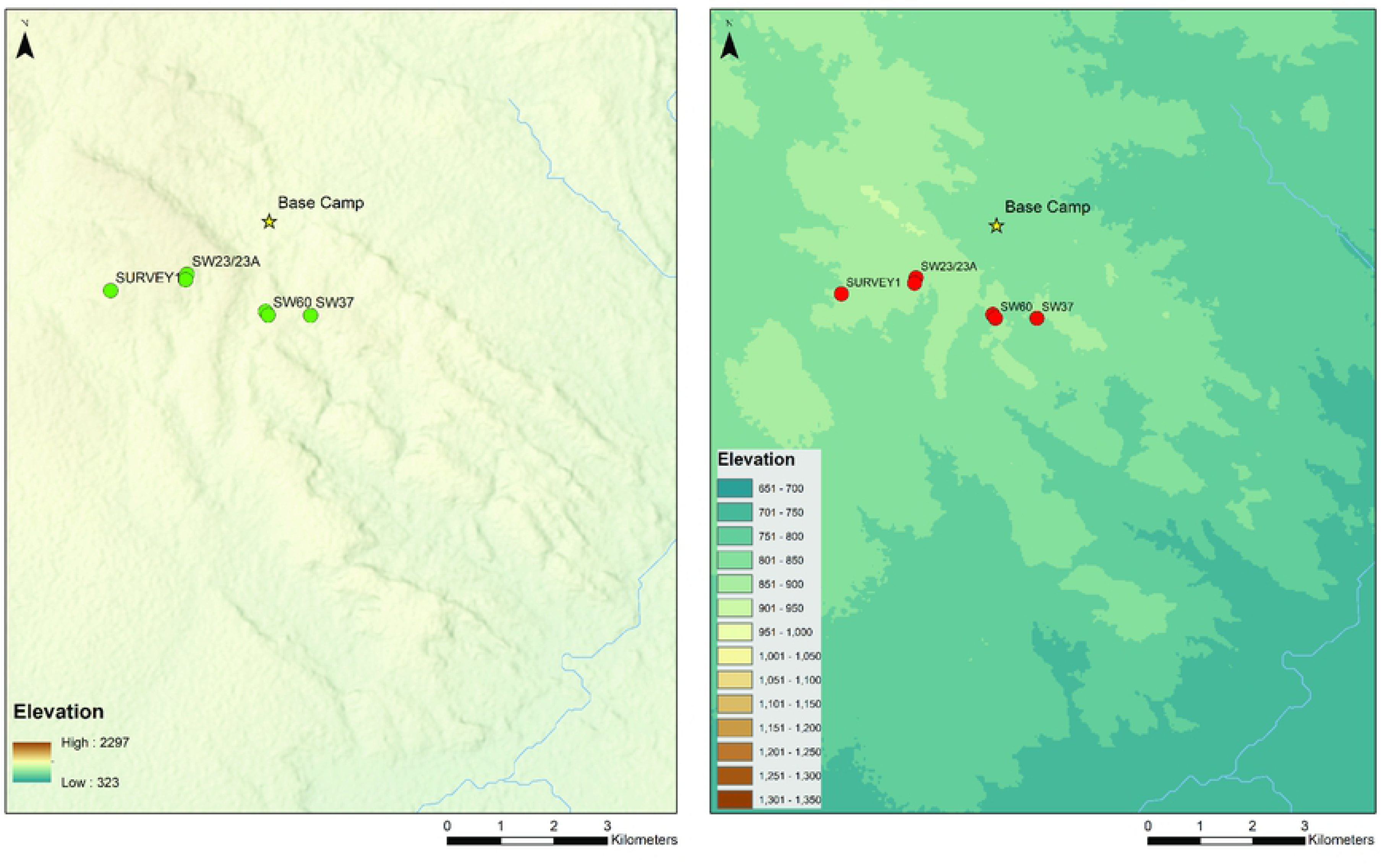
Phase 2 stratified archaeological survey localities (2016, 2019)

Phase 2, stratified archaeological surveys were conducted at five localities in 2016 and 2019 by teams of 2-4 trained operators walking parallel transects at ca. 8 m intervals, recording all surface finds. Survey tracks and artefacts were recorded using hand-held Garmin © GPS devices.

Spatial data from the GPS units, including survey tracks and waypoints, were processed in the field using Garmin ©Basecamp software and later exported to a geographic information system, or GIS (ESRI ©ArcMap 10.8) for post-processing and analysis. The GIS includes provisions for pointfiles (locations/localities/artefacts), feature classes (polylines for survey tracks, polygons for surveyed areas) and raster map layers (base maps). Base maps incorporated in the GIS include a 30-m SRTM DEM, background imagery (ESRI web-hosted layer), rasterised topographic maps (series Z741, D.O.S. 424, sheets 1033C1 & 1033C3, 1:50,000 scale), geological maps (degree sheet 1033 SW & SE Muyobe & Luwumbe, Geological Survey Department ZGS, 1:100,000 scale) (27), landcover from Sentinel 2 data processed by the Zambian ESA-CCI (European Space Agency–Climate Change Initiative) Land Cover (LC) team, and rainfall erosivity (GlobalR) from the European Soil Data centre (ESDAC). Topographically derived layers (slope, hydrology) were produced using ArcMap tools. Regional-scale administrative boundaries, hydrology, rainfall and erosivity data were sourced from HydroAtlas_Zambia v. 1.0 (16).

Phase 1 surveys were used to provide a broad picture of the regional geological and geomorphological context within which artefact scatters occur. Descriptive statistics were produced using ggplot and regression analyses were performed using the AICcmodavg package in R (33). For Phase 2, survey tracks were merged and buffered using a 2-m buffer and superimposed on a 1x1 meter grid in order to calculate the spatial extent of the surveys and artefact densities using standard arcMAP tools. A kernel density function was used to identify artefact clusters within the survey areas. Statistical summaries were produced for the survey areas and larger artefact clusters using zonal statistics.

Techno-typological identification of lithic artefacts was conducted in the field in 2016. In 2019 artefacts were georeferenced and collected for analysis at the University of Lusaka, including typological identification and the recording of detailed morphological, metrical and taphonomic attributes following Barham (34) and Clark et al (35). Excavated material and surface artefacts collected from SW23B in 2019 are reported separately, together with the results of archaeological excavations conducted at the locality (Bisson et al. forthcoming).

### Paleoenvironmental analyses

A long sedimentary sequence exposed by gullying at SW23, between SW23A and SW23B. was described in the field. A total of 11 sediment samples were collected from the profile of a test trench cut into the slope above SW23B for geoarchaeological and paleoecological analyses, including grain-size analysis, palynology and phytolith analysis; a further six block samples and five loose samples were collected for micromorphological analysis (SI-2), five additional sediment samples were collected for OSL dating (Fig. 7). Grain-size analysis was undertaken by first treating each sample overnight with 10% hexametaphosphate solution to disaggregate the sediment, followed by agitation using a vortex mixer for one minute, and then applying a sonicator to break-up any remaining aggregates, before analysis using a Betatek Laser Particle Size Analyzer. Only the grain-size distribution of the <2mm fraction was assessed as this is the maximum size the analyzer permits. Bulk elemental analysis was also undertaken on each sample using an Olympus Vanta^TM^ handheld XRF analyzer to obtain concentrations of a suite of elements, including Al, Si, P, Cl, Ca, Ti, Mn, Fe, Ni, Cu, Zn, Rb, Sr, Y, Zr, Nb, Pb, and Th.

**Fig 7.**
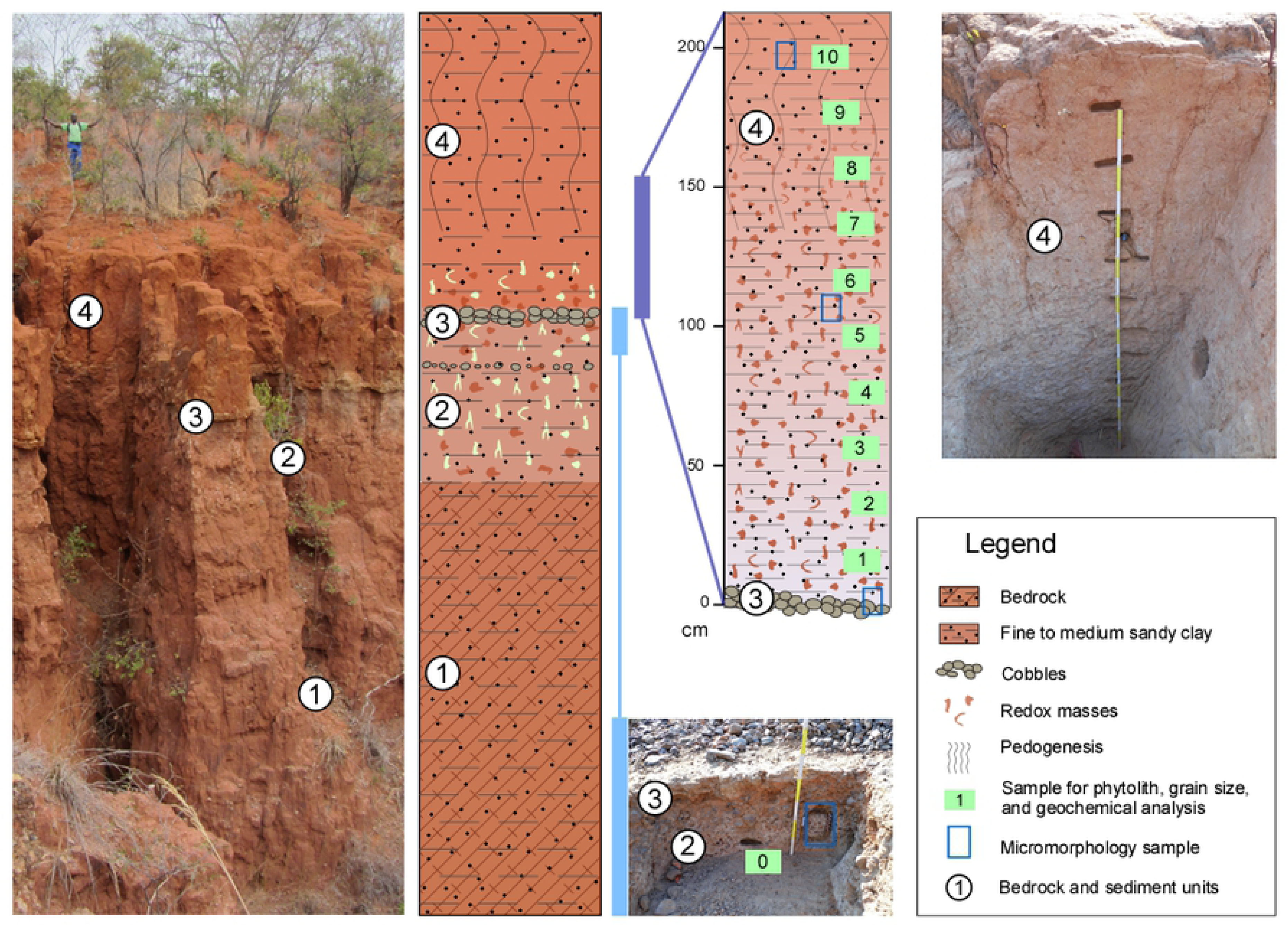
Sediment column at SW23. Exposure between SW23A and B showing location of samples collected for sedimentological and environmental analyses.

Standard pollen processing protocols were used (36), to concentrate pollen and spores, but a number of scans of slides at both 200x and 400x magnification indicated that the samples were sterile. Phytolith analyses were conducted on 1.5 to 3 g of sediment for each sample. The extraction protocol is based on (37) and details are provided in SI. Phytoliths were described according to their three-dimensional shape and classified following the international code of phytolith nomenclature (38, 39) and the classification schemes described in SI.

## Results

### Phase 1: regional context

The piedmont surface is covered by a thin, discontinuous mantle of cobbles, flakes and isolated artefacts, extending into river valleys. Cobble concentrations tend to occur on sloping ground on the edge of the piedmont zone and in riverbank deposits and cuttings. A total of 63 locations were explored during Phase 1, on the upper piedmont surface within the Luwumbu catchment and part of the Viziba. Survey locations are associated with the Upper Grit Formation (T, N = 24), the Ntawere Formation (S2, N = 22) and Madubisa Mudstone formations (R3, R2, N = 15) (Fig 5b). Concentrations of lithic artefacts were encountered in 8 localities (12% of the total sample) in the piedmont zone. Artefacts co-occur with fossil wood in two locations (3% of the sample) and with Triassic vertebrate remains in a location associated with the Ntawere Formation (S2) (SI_1).

Exploratory statistics show that with one exception, Phase 1 localities with stone tool concentrations occur on the Upper Grit Formation above 800 meters asl, on relatively sloping ground on the edge of the upper piedmont zone. Globally, lithic scatters occur in areas with relatively low soil erosivity, which reflects the generally high erosivity values in the river valleys where cultivated fields occur (Fig. 8). Within the piedmont zone, lithic scatters occur in areas with relatively high soil erosivity. They also tend to occur in zones categorized as either grassland or shrubland, according to the 2016 Sentinel 2 landcover data, although field observations suggest they occur in thin miombo forest. Predictor variables tested using regression analysis (GLM) include elevation, slope and soil erosivity. None of the candidate predictors proved significant, with the exception of soil erosivity, on a regional scale.

**Figure 8:**
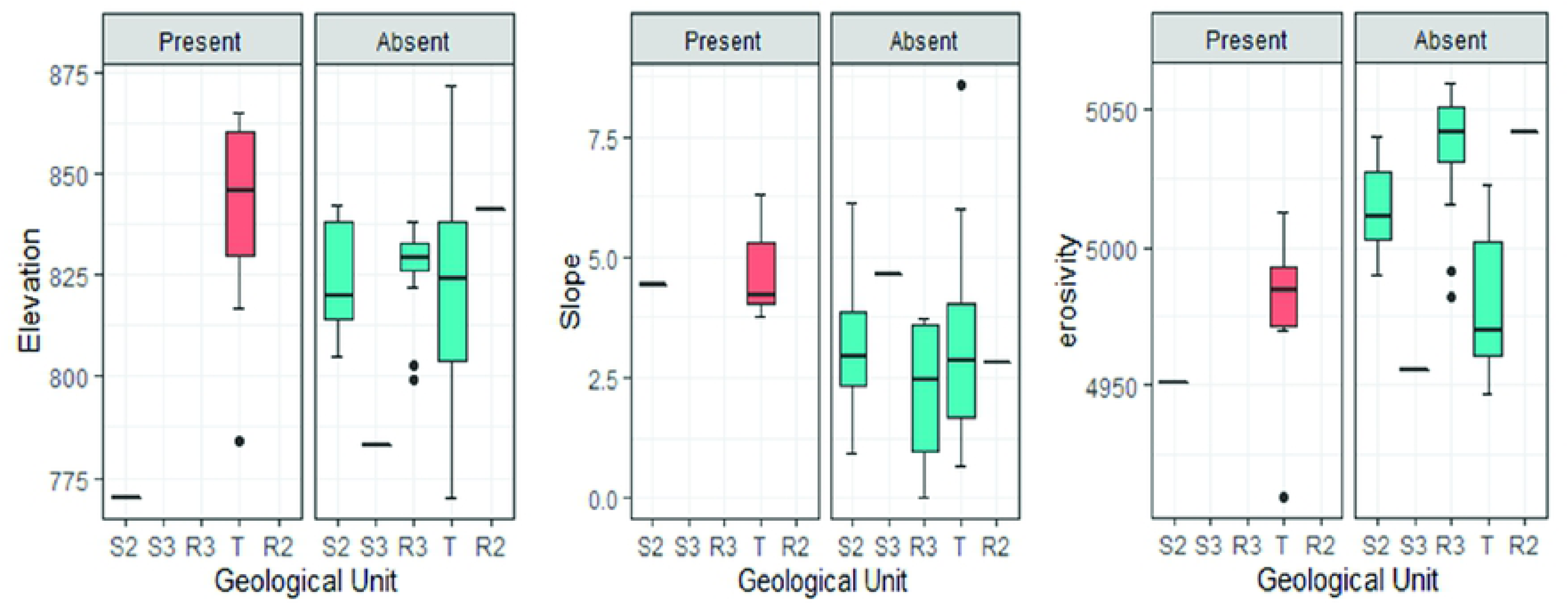
Exploratory statistics: Phase 1 localities. The presence/absence of lithic artefacts as a function of elevation, slope and erositivity by geological unit.

### Phase 2: stratified, archaeological surveys

The localities selected for Phase 2 archaeological surveys consist of relatively dense concentrations of cobbles and lithic artefacts situated on the margins of the piedmont zone, on strong to moderately sloping ground at elevations ranging from 850 to 900 m asl, in areas characterised by slightly higher erosion potential. Four of these localities (SW23 A & B, SW37, SW60) are situated at the headwaters of the Lutete and the Nkande rivers, two seasonally active watercourses that form a sub-catchment draining into the Luwumbu River. A fifth locality is situated further west on the upper piedmont surface (SURVEY1), near the headwaters of the Viziba River, which drains into the Luangwa.

Two of the survey localities, SW23 A & B, are located on adjacent spurs separated by a deep gully (ca. 20m deep). Dense concentrations of artefacts are present at both locations (Table 1) which form part of Phase 2. Phase 1 exploratory surveys conducted in 2019 revealed the presence of cobbles, flakes and isolated heavy tools on an adjacent, less eroded feature with relatively more vegetation cover.

**Table 1.**

| PHASE 2 LOCALITIES | SURVEY AREA (m<sup>2</sup>) | POINTS N | CLUSTERS N | DENSITY Max (m<sup>2</sup>) | DENSITY Mean (m<sup>2</sup>) | % SLOPE | SLOPE |
| --- | --- | --- | --- | --- | --- | --- | --- |
| SW23A | 2280 | 402 | 2 | 14 | 0.18 | 10 | Moderate |
| SW23B | 792 | 1661 | 7-8 | 28 | 2.09 | 7 | Gentle |
| SW37 | 1029 | 176 | 3 | 8 | 0.17 | 9-14 | Moderate |
| SW60_2016 | 857 | 79 | 4 | 6 | 0.09 | 4 | V. Gentle |
| SW60_2019* | 172 | 53 | 2 | 6 | 0.31 | 9-13 | Moderate |
| SURVEY1 | 55 | 14 | 1 | 3 | 0.25 | 10 | Moderate |

*\*survey of an adjacent zone to the zones surveyed in 2016*

At several locations on the landscape, including but not limited to SW23A, SW23B and SW60, large clasts protect softer, underlying sediments from erosion. Annual surface runoff values in the study region (including the Luwumbu and Viziba catchments) are relatively high, ranging from 172 to 189 mm, and peak in the rainy season (16). As the surrounding fine fraction of sediment is removed from around the large clasts by sheet wash, pedicles ca. 20-30 cm high form capped by cobbles, large flakes, and lithic artefacts (Fig. 9).

**Figure 9:**
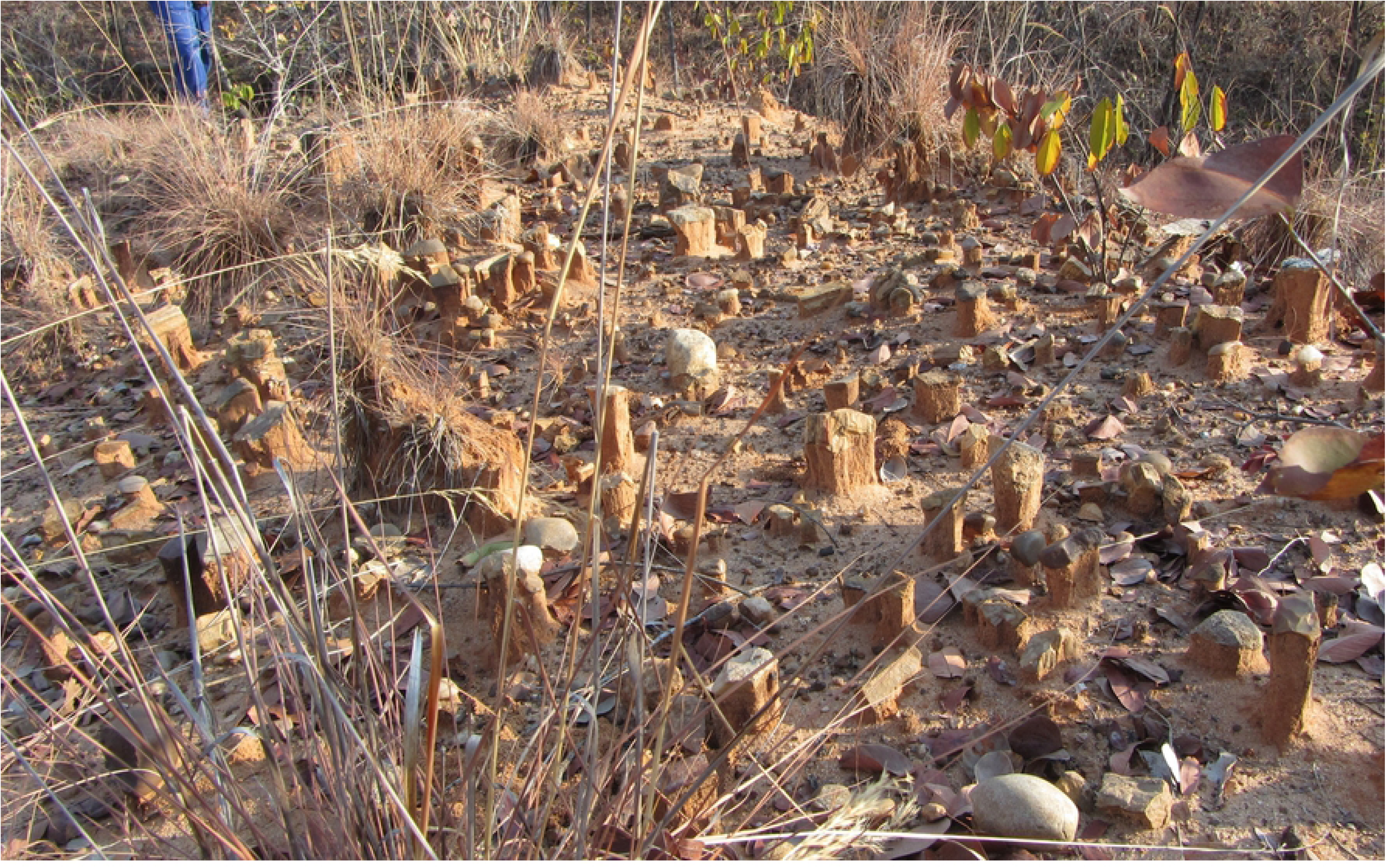
Sheet-wash action. Pedicles under large clasts and artefacts formed as the fine fraction is removed by sheet-wash (here in the vicinity of SW60).

**Figure 10:**
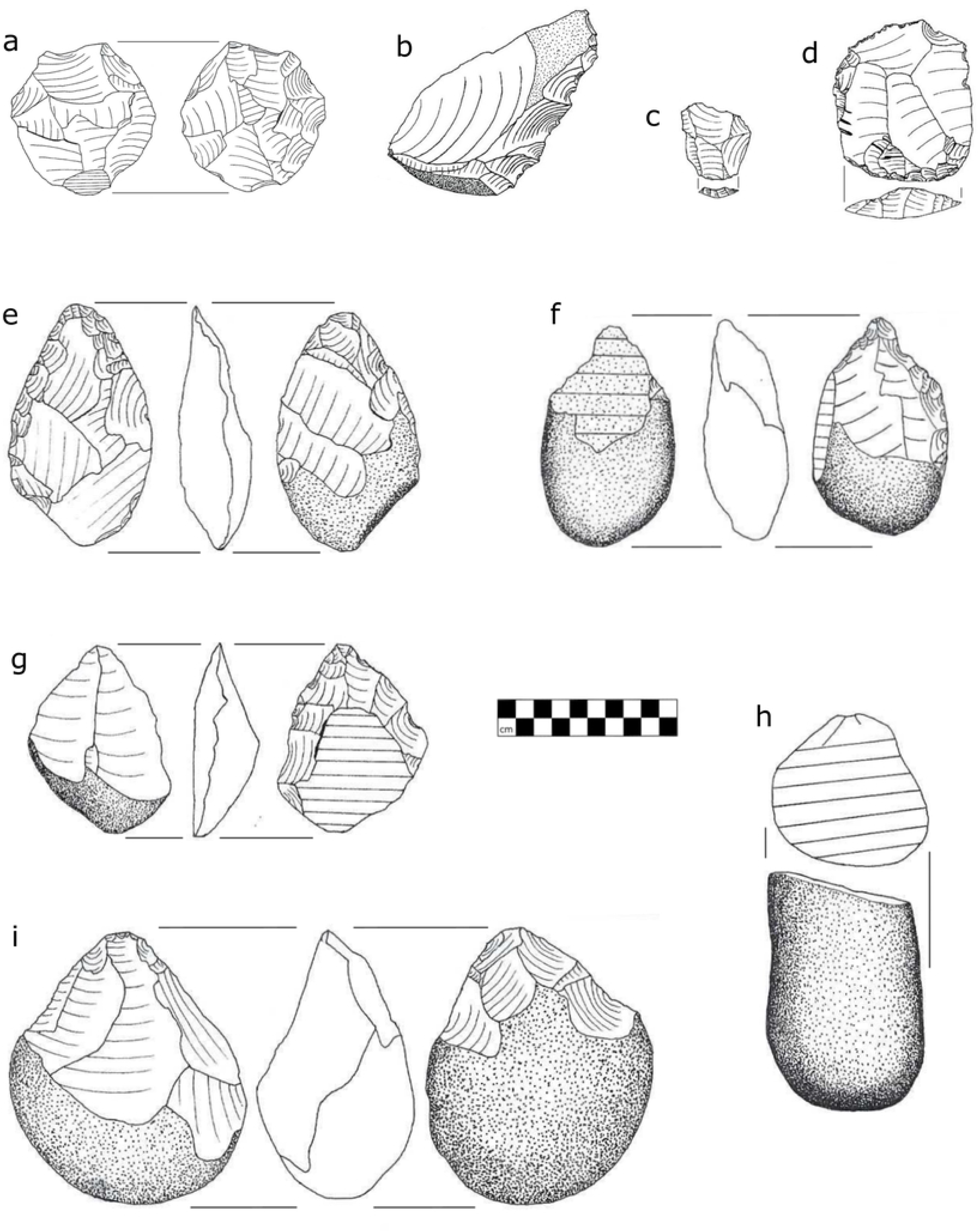
Artefacts from Sitwe Localities. a, Radial core (SW23A); b, Denticulated side-struck flake (SW23B); c, Levallois flake (SW23B); d, Levallois cleaver flake (SW23A); e, Handaxe (SW23A); f, Pick; g, Unifacial handaxe on a split cobble spall (SW60); h, Elongated split cobble (SW23A); i, Core-axe (SW23A). Scale in cm.

### Lithic Analysis

Archaeological material from four Phase 2 survey localities (SW23A, SW23B, SW60 and SW37) is presented here (Table 2). The fifth locality, SURVEY1, represents a much smaller, low-density scatter dominated by flakes, similar to the “between the patches” scatters described by Foley (40); discovered in 2019, it has yet to be studied. Two Phase 1 survey localities (MO1, SW22) are also described.

**Table 2.**
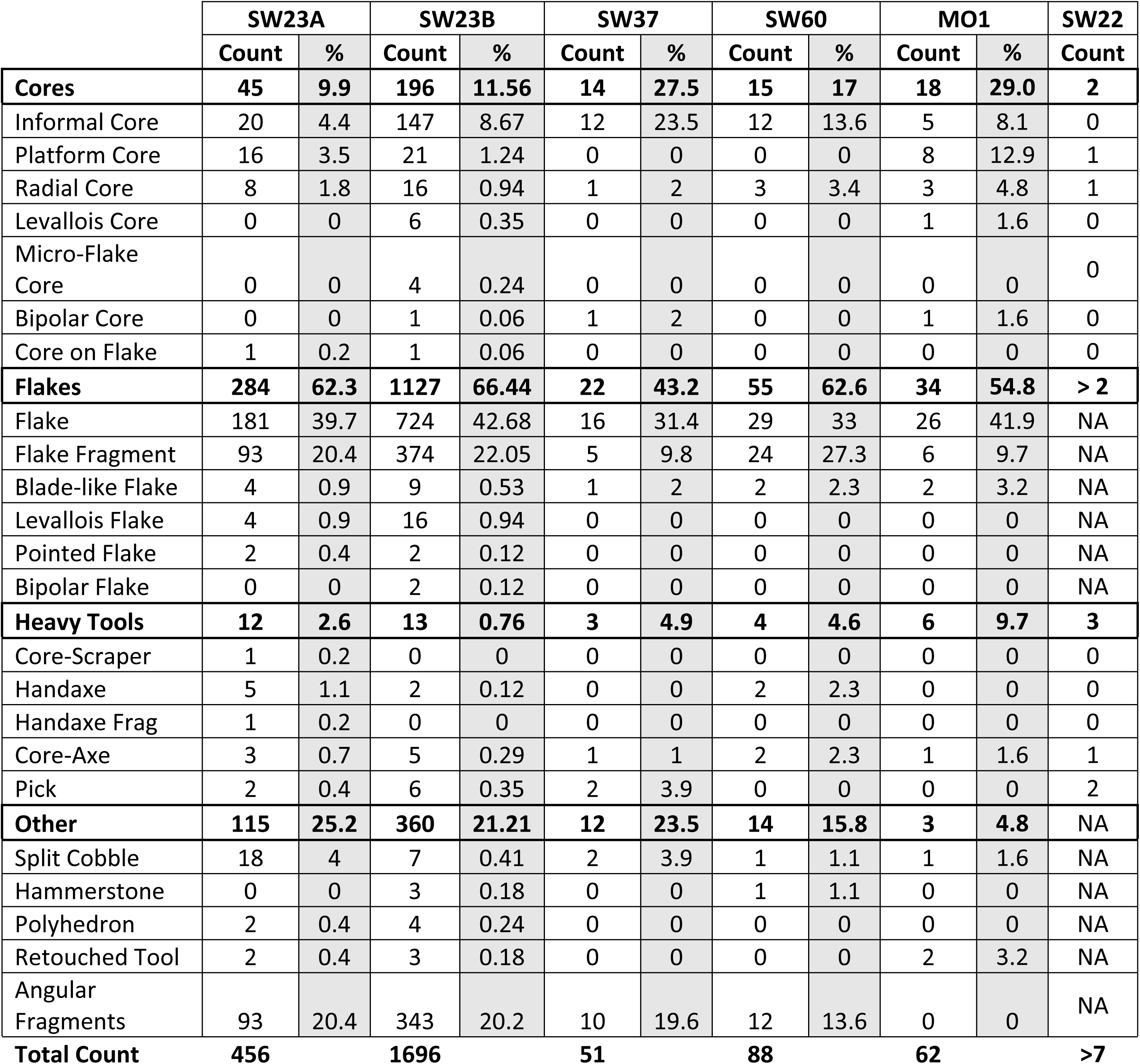

|  | SW23A |  | SW23B |  | SW37 |  | SW60 |  | MO1 |  | SW22 |
| --- | --- | --- | --- | --- | --- | --- | --- | --- | --- | --- | --- |
|  | Count | % | Count | % | Count | % | Count | % | Count | % | Count |
| **Cores** | **45** | **9.9** | **196** | **11.56** | **14** | **27.5** | **15** | **17** | **18** | **29.0** | **2** |
| Informal Core | 20 | 4.4 | 147 | 8.67 | 12 | 23.5 | 12 | 13.6 | 5 | 8.1 | 0 |
| Platform Core | 16 | 3.5 | 21 | 1.24 | 0 | 0 | 0 | 0 | 8 | 12.9 | 1 |
| Radial Core | 8 | 1.8 | 16 | 0.94 | 1 | 2 | 3 | 3.4 | 3 | 4.8 | 1 |
| Levallois Core | 0 | 0 | 6 | 0.35 | 0 | 0 | 0 | 0 | 1 | 1.6 | 0 |
| Micro-Flake Core | 0 | 0 | 4 | 0.24 | 0 | 0 | 0 | 0 | 0 | 0 | 0 |
| Bipolar Core | 0 | 0 | 1 | 0.06 | 1 | 2 | 0 | 0 | 1 | 1.6 | 0 |
| Core on Flake | 1 | 0.2 | 1 | 0.06 | 0 | 0 | 0 | 0 | 0 | 0 | 0 |
| **Flakes** | **284** | **62.3** | **1127** | **66.44** | **22** | **43.2** | **55** | **62.6** | **34** | **54.8** | **> 2** |
| Flake | 181 | 39.7 | 724 | 42.68 | 16 | 31.4 | 29 | 33 | 26 | 41.9 | NA |
| Flake Fragment | 93 | 20.4 | 374 | 22.05 | 5 | 9.8 | 24 | 27.3 | 6 | 9.7 | NA |
| Blade-like Flake | 4 | 0.9 | 9 | 0.53 | 1 | 2 | 2 | 2.3 | 2 | 3.2 | NA |
| Levallois Flake | 4 | 0.9 | 16 | 0.94 | 0 | 0 | 0 | 0 | 0 | 0 | NA |
| Pointed Flake | 2 | 0.4 | 2 | 0.12 | 0 | 0 | 0 | 0 | 0 | 0 | NA |
| Bipolar Flake | 0 | 0 | 2 | 0.12 | 0 | 0 | 0 | 0 | 0 | 0 | NA |
| **Heavy Tools** | **12** | **2.6** | **13** | **0.76** | **3** | **4.9** | **4** | **4.6** | **6** | **9.7** | **3** |
| Core-Scraper | 1 | 0.2 | 0 | 0 | 0 | 0 | 0 | 0 | 0 | 0 | 0 |
| Handaxe | 5 | 1.1 | 2 | 0.12 | 0 | 0 | 2 | 2.3 | 0 | 0 | 0 |
| Handaxe Frag | 1 | 0.2 | 0 | 0 | 0 | 0 | 0 | 0 | 0 | 0 | 0 |
| Core-Axe | 3 | 0.7 | 5 | 0.29 | 1 | 1 | 2 | 2.3 | 1 | 1.6 | 1 |
| Pick | 2 | 0.4 | 6 | 0.35 | 2 | 3.9 | 0 | 0 | 0 | 0 | 2 |
| **Other** | **115** | **25.2** | **360** | **21.21** | **12** | **23.5** | **14** | **15.8** | **3** | **4.8** | **NA** |
| Split Cobble | 18 | 4 | 7 | 0.41 | 2 | 3.9 | 1 | 1.1 | 1 | 1.6 | NA |
| Hammerstone | 0 | 0 | 3 | 0.18 | 0 | 0 | 1 | 1.1 | 0 | 0 | NA |
| Polyhedron | 2 | 0.4 | 4 | 0.24 | 0 | 0 | 0 | 0 | 0 | 0 | NA |
| Retouched Tool | 2 | 0.4 | 3 | 0.18 | 0 | 0 | 0 | 0 | 2 | 3.2 | NA |
| Angular Fragments | 93 | 20.4 | 343 | 20.2 | 10 | 19.6 | 12 | 13.6 | 0 | 0 | NA |
| **Total Count** | **456** |  | **1696** |  | **51** |  | **88** |  | **62** |  | **>7** |

Globally, cores account for 12 to 27% of the lithics recovered from Phase 2 survey localities, and flakes and flake fragments represent 43 to 66% of the assemblages (Table 2). Most of the lithic material is not diagnostic, consisting of angular fragments, unmodified flakes, flake fragments, and informal cores. Retouched tools are rare and include a single scraper, notches and a longitudinally split Levallois flake denticulated on the interior face of its sharp lateral edge. Heavy tools such as bifaces, core-axes, handaxes and picks, and other diagnostic tool types are present but poorly represented. Most of the lithic material encountered is the product of basic flaking activities using free-hand, hard hammer percussion. Detached pieces are dominated by partially cortical flakes produced by hard hammer percussion from informal or simple platform cores, the most common core type. The diversity of core types increases with sample size and includes Levallois, blade, bipolar and radial cores.

Split cobbles occur in small numbers in all four Phase 2 localities and are the most common heavy-duty artefact. They appear to be deliberate products rather than tested and discarded raw material (14, 41, 42) or initial stages in the production of single platform or radial cores (43, 44). Instead, elongated cobbles were split, possibly by bi-polar percussion, perpendicular or slightly oblique to their long axis, creating an elongated piece with a flat facet on one end. Large cutting tools (LCTs) including an unusual unifacial form of handaxe made on a prepared split cobble (Fig 8c) are present at all four localities, in addition to two Phase 1 localities whose lithics have been described (Table 2). Handaxe forms are primarily elongated ovate or lanceolate and most have unflaked butts. One ovate specimen was made on a giant side-struck flake and expertly thinned with soft-hammer percussion. An unusual handaxe type found at both SW23A and SW60 was made by unifacially retouching the interior face of a giant spall split from an elongated cobble or boulder. Prior to splitting the cobble, two large flakes were struck unifacially from one end of the support creating a ridge. The cobble was split so that this ridge terminated at one end of the resulting spall and that was the end that was retouched to create a thin pointed handaxe tip.

The lithic material displays a range of depositional and post-depositional alteration. Surface condition ranges from fresh and sharp to abraded and rolled, patinated and unpatinated. The percent of flakes with some cortex ranges from just over 56% (SW23B) to 91% (SW37). The most common raw materials used for knapping are quartzite and white (crypto-crystalline) quartz cobbles, although fossil wood is also occasionally exploited (most notably at MO1; Table 2) doubtless because natural flaws make it unreliable for flaking despite its’ fine grain.

### Geoarchaeological analyses

#### Sediment sequence

The deeply incised gully system between SW23A and B affords an overview of the local depositional sequence (Fig.7) which is composed of four main units. The sediment sequence is undated as the OSL samples were damaged in transit.

Unit 1: Sandstone bedrock of the Upper Grit formation (Upper Karoo) is exposed on the gulley bottom, underlying the sedimentary units. The sandstone is fine-grained and exhibits a dark red color which has formed after exposure due to oxidation and weathering, as observed in the field and in micromorphological thin sections (Fig SI_2-1:i-l). The color is prevalent on the outside of the rock and along root disturbances. Lower down, the sandstone is pale, clay-rich quartz with minor contribution of feldspar, based on observations in thin section.

Unit 2: The deposit overlying the sandstone bedrock consists of ca. 1,5 m of fine sandy clay similar in composition to the sandstone in terms of mineralogy and grain size. The deposit contains reworked sandstone fragments (1-5 cm) of the Upper Grit (1), recognizable by their red color but difficult to differentiate from *in situ* redox masses (mottles), which are common in lower parts of the soil profile affected by groundwater (Fig. SI_2-1: d,e,g). A first surge of higher-energy deposition is represented by a locally appearing distinct pebble stringer running 30 cm below the top of this unit, and elsewhere by a general increase in coarse pebbles and cobbles in the top 30 cm, with common feldspars.

Unit 3: This unit consists of a ca. 15-40 cm bed of imbricated cobbles (15-20 cm in diameter) of grey quartzite with few feldspars or other rock types (Fig. SI_2-1:f). In the exposed gully sections in the vicinity of SW23A and SW23B, this cobble layer can be followed and observed as an undulating, wavy unit that is locally cemented by iron-manganese concretions (Fig. SI_2-1:e). The cobble unit outcrops in some areas on the terrace of SW23B, especially near the edges.

Unit 4: The cobbles are overlain by Unit 4, which starts at its base with bedded medium gravel (ca. 10 cm) before transitioning to fine to medium sandy clay. This upper sedimentary unit is three to five meters thick at the gully exposure, depending on the extent of erosion at the top of the sequence. The sediments from the test trench cut into the slope above SW23B show limited soil development as they had been deeply buried until comparatively recently. Grain size analysis revealed fine intact stratification of the sandy clay at depth, with slight variations observed in the sand fraction. Inclusions of very coarse sand (1-2 mm) were only present in one sample (PP5) from the T1 column (Fig.7) and the prevalence of fine sediments appears to indicate a persistent, low-energy depositional environment.

Lateritic soil formation has overprinted Unit 4. While pedogenesis is relatively recent and limited at the sampled test pit (T1) due to its recent exposure (Fig. SI_2-1:a-c), in the large gully exposures it has visibly impacted several meters of sediment and a thick (> 2 m) upper zone, or biomantle (45, 46) of homogenized pinkish red sediment is observed, which transitions into a mottled zone 1-2 meters above the cobbles of Unit 3 or lower, depending on the burial depth and progression of erosion of Unit 4. Between the mottles and underneath, the deposit is pale in color reflecting the non-oxidized, reduced color of clays and sand in the pallid zone of a groundwater-impacted profile (47). The elemental geochemical results (SI_3) show that the Al/Si ratio is elevated toward the base of the section (PP2 and PP3). This ratio has been used as a proxy for the relative proportion of clays to quartz sand, and thus may reflect enhanced chemical weathering (48, 49), although the grain size results (SI_3) do not show a higher proportion of clay particles at this level.

#### Palaeoenvironmental analyses

Standard pollen processing protocols (36) yielded no results, but phytoliths were found in all the samples except for PP3. The number of phytoliths counted varies from 302 to 431 and is sufficient for identification and the computation of indices.

Phytolith assemblages in all eleven sample are dominated by globular granulate morphotypes indicative of woody dicotyledons, representing more than 50% of the assemblage (SI_4). The tree cover index, D/P ratio, was > 2 for all the samples, which corresponds to close formations such as Miombo woodlands or forests. Grass phytoliths were found in the samples and dominated by the lobate morphotypes (bilobate and cross) corresponding to the Panicoideae subfamily. These C4 grasses, present in all the samples, suggest that the environment was probably a woodland. The Iph ratio (aridity index) indicated changes in grass composition and climatic conditions. Samples PP-10, 9, 7, 5 and 2 have Iph ratios < 20%, which indicates high available soil moisture conditions and potentially a more humid climate. Conversely, samples PP8, 6 and 4 have Iph ratios > 40% suggesting more arid conditions. While PP0 and PP1 lie within the intermediate ratio values, it is possible that PP0 corresponds to more xeric conditions (39%), and PP1 to more humid ones (20%). It should be noted, however, that less than 30 phytoliths used to compute the Iph ratios for PP4, 5, 6 and 10. This should not affect the contrast between wet and dry conditions but makes further interpretation impossible.

## Discussion

The results of our fieldwork in the Luwumbu sub-basin and adjacent regions confirm that the piedmont zone in northeastern Luangwa retains a record of hominin occupation that potentially spans the Early to Late Stone Age. The survey localities reported above consist of concentrations of cobbles and lithic artefacts situated on the upper piedmont zone at elevations between 850-900 m asl. They are readily distinguishable from the relatively thin and discontinuous mantle of cobbles, flakes and artefacts that covers the piedmont surface based on their relative density.

The spatial and temporal integrity of the surface deposits investigated above calls for a landscape-based, geoarchaeological approach (50). Long-term processes, operating at large spatial scales, are the only possible explanation for the co-occurrence of Stone Age artefacts, fossil wood and/or fossil vertebrates on the contemporary piedmont surface, and their co-occurrence in some of the survey localities (e.g., SW37). Reconstructing the sequence of geomorphological events that resulted in the formation of the piedmont and subsequent patterns of sediment deposition, therefore, is critical to understanding the presence of archaeological material in surface deposits and assessing the probability that *in situ* archaeological deposits may be preserved in the region.

Our field observations and previous fieldwork in the region (2, 19) suggest that the Upper Grit formation in the northern Luangwa Valley formed a topographically complex surface that was subsequently partially covered by piedmont fans flowing from the Mafinga Hills and the Makutu Range. Based on the size of the clasts contained in the fanglomerates [*Neil Tabor, pers. comm. 2021*] the fans likely formed braided streams rather than debris flows, such as the ones documented in the foothills of eastern Luangwa, near Chipata (8). The deposition and coalescence of the piedmont fans, which partially infilled low-lying areas on the Upper Grit surface, and subsequent erosion of outcropping Upper Grit sediments, resulted in the formation of an undulating piedmont surface. A Quaternary sediment package of variable depth, consisting primarily of alluvium, was then deposited on this surface. Finally, large-scale cycles of aridification, which have shaped the climate of Central Africa since the Miocene, will have resulted in alternating cycles of sediment deposition and incision of the piedmont surface. Macro- and micromorphological observations of sediment samples recovered from SW23 are consistent with this reconstruction of events, which suggests that the piedmont zone has a long and complex history.

The long sediment sequence exposed by gullying between SW23A and SW23B, on the edge of the upper piedmont zone, includes a more than 3m thick sediment package overlying an imbricated cobble level and ∼1.5 m of underlying deposits resting on the Upper Grit. The upper sediment package could correspond to a sequence of Quaternary sediments deposited on the piedmont surface. For the moment we lack the necessary chronological controls to establish whether this is the case, nor can we determine when the cobble level was deposited, which would provide clues as to the timing of piedmont formation.

Geoarchaeological analysis of the sediment sequence at SW23 indicates that initial formation of the piedmont zone involved a phase of high-energy deposition, possibly an alluvial fan, a braided river system or meandering stream (Unit 3). The cobble bed representing this episode is the most distinct feature of SW23 and was not clearly identified at the other survey localities (SW37, SW60, SURVEY1). The absence of pedogenic development in the deposits underlying the cobble level (Unit 2) and similarity to the sediments overlying the cobbles precludes a long-term hiatus in sedimentation. Continued deposition above the cobble bed is characterised by fine sandy deposits, with slight variations in grain size representing long-term, low-energy alluvial deposition (Unit 4) after initial piedmont formation. The erodibility and ubiquity of the sandstone bedrock in the area generated a ready source of sediment in the past and similar processes of sandstone weathering and re-deposition are ongoing in the region. The sandy clays under- and overlying the cobble layer are in part composed of the weathering product of the Upper Grit Formation, the clay results from the weathering of feldspars.

The structural terrace of SW23B and adjacent features represents an erosional landform. Triggered by monsoonal rainfall between November and March, erosion by sheetwash [FIG] and gullying are potentially the driving factors behind the formation of the terrace by removal of sediments of Unit 4. The removal of fine sediment has reached the level of the cobbles at the edges of the terrace, while ca. 70 cm of Unit 4 sediment was observed above the cobbles in central parts of the terrace. A similar process is underway at SW23A. This may explain why the archaeological deposits from these localities, and others in the region, are typologically mixed. At SW23B, however, field observations of sediment exposures and archaeological excavations (reported elsewhere) also reveal the presence of some archaeological material, including flakes and informal cores, in the undated sediment package overlying the Upper Grit, hinting at the untapped archaeological potential of the site.

Fossil vertebrates, fossil wood and archaeological material all occur on the contemporary piedmont surface above ∼850 m asl but are, for the most part, spatially segregated indicating that intact sequences of Quaternary sediment exist, separated by outcrops of Upper Grit. Older sediment exposures occur when overlying Quaternary sequences have been stripped away by erosion, e.g., on the lower margins of the piedmont where paleontological fieldwork has concentrated.

The cobbles that mantle the contemporary piedmont surface could derive from the Makutu Range, which consists mainly of schists and quartzites overlying basal granite (27–29). They may also have originated in the more distant Mafinga Hills. Their presence could reflect relatively recent, episodic flooding events and/or the gradual exposure of previously buried clasts through erosion. Concentrations of large clasts, including cobbles and lithic artefacts, likely represent time-averaged lag deposits formed by erosion and their spatial distribution reflects the underlying topology of the piedmont surface. They also represent raw material sources that were actively exploited for toolmaking throughout the Stone Age in the study region.

Typologically, archaeological material from the Luwumbu sub-basin spans the Early and Middle Stone Age, confirming that hominins occupied northeastern Luangwa more or less continuously after initial piedmont formation. The lithic assemblages (see: Table 2) are lag deposits dominated by flakes and minimally reduced cores; the larger assemblages (e.g., SW23A, SW23B) are more typologically diverse but the bulk of the evidence points to occupation during the MSA or early MSA (EMSA). Further analysis is complicated by the lack of chronological control and the fact that the MSA in Africa is highly variable, both temporally and spatially. Archaeological fieldwork conducted in the southern sub-basin provides a basis for comparison and clues as to the possible age of the deposits from which the surface assemblages in northeastern Luangwa are derived.

Extensive geoarchaeological fieldwork in central Luangwa, in the southern sub-basin along a transect of the floodplain from the Nchindeni Hills to the Muchinga escarpment, was undertaken as part of the Past Peoples and Environments Project (12, 13). In addition to documenting secondary archaeological deposits in the floodplain, a series of coarsely stratified deposits containing Stone Age tools capping a dissected pediment located ∼20 m above the modern floodplain, on the right bank of the Luangwa River, was investigated. These perched gravels are thought to result from the active dissection of alluvial fan deposits linked to climate-driven flash floods occurring at some point during the Middle or Late Pleistocene (13). Broad patterns in lithic production are observable in the materials excavated from the hilltop localities (12). Retouched tools include types ascribable to the LSA in the upper levels, while heavy-duty tools (e.g., picks) attributed to the ESA and early MSA occur further down the sequence. Similarly, core samples show broad trends from top to bottom: centripetal, flaked cores occur in the upper sequences; radial and disc cores occur further down and split and flaked cobbles occur towards the base of the sequence. Finally, diachronic patterns in raw material use correlate with differences in the size of flakes produced, with larger flakes in the lower levels correlated with increased use of quartzite (Op. Cit.).

At Manzi, a Stone Age locality situated near the confluence of the Manzi and the Luangwa rivers just south of the transect studied by Colton, a discontinuous sequence of fluvial and colluvial deposits containing cobble layers and small numbers of artefacts in secondary context, unconformably overlies Karoo sediments (6). Six sequences of fine sediments demarcated by cobble layers were identified, from top to bottom: CL1 to CL6 (6:Fig.3). CL1 to CL3 (the Upper Unit) are separated from CL5 and CL6 (the Lower Unit) by an unconformity (CL4). The fluvial depositional context means that artefacts are in secondary context; abrasion patterns suggest CL6, and especially CL5 were subjected to relatively more transport damage. Cosmogenic nuclide dating of the Manzi sequence was unsuccessful but indicates that the Lower Unit was considerably eroded before being buried by overlying, younger deposits. Techno-typological attributions are complicated, as the authors point out, due to the paucity of retouched tools and the difficulty inherent in attributing basic reduction techniques, associated with Mode 1, to either the ESA or the MSA. The Lower Unit is dominated by Mode 1, lacks LCTs but contains centripetally flaked split cobbles (disc cores) and multiple platform cores. The Upper Unit contains prepared cores associated with Mode 3 (MSA). Paleomagnetic data for the Lower Unit points to a transition from a normal subchron to the reversed polarity of the Matuyama chron (2.14 - 0.78 Ma) and Barham and colleagues suggest a date no older than 1 Ma. The age of CL3 is unknown but the uppermost layers (CL1 and CL2) accumulated during the Brunhes polarity and isothermal dating techniques indicate an age of ca. 78 ka for CL2.

In summary, lithic assemblages from central and northeastern Luangwa are dominated by simple flake production, complicating chronological attributions. Relative chronologies have been established in the central region (above) indicating a hominin presence from at least circa 1 Ma associated with a simple core reduction technology (Mode 1), to more recent, MSA (Mode 3) and LSA (Mode 5) occupations of the valley during the Middle/Late Pleistocene.

Geomorphological evidence points to periodic pulses of climate-driven, high-energy deposition in the floodplain and in the foothills of the Luangwa valley. Their precise chronology is as yet undetermined, but they washed good-quality raw material down from nearby hills, forming successive cobble deposits that were eventually buried and then locally exposed by incision. The availability of these reliable sources of raw material in the Luangwa valley could explain the prevalence of simple cores and flakes in the archaeological assemblages. The earliest hominin occupation of the piedmont zone in northeastern Luangwa could be roughly contemporaneous with the lower levels at Manzi, for which an *ante quem* date of ∼1 Ma is suggested, but the presence of LCTs suggests the ESA or early MSA (EMSA). LSA components in the study region are very rare, a pattern that differs somewhat from observations in nearby Malawi (51) and Mozambique (52), indicating that the timeframe during which the deposits from which the lithic material in the piedmont zone in northeast Luangwa derives may be somewhat older.

### Lithic provisioning

The information gathered from the lithic scatters studied for this project does not include a small fraction, which would have allowed us to draw conclusions about hominin mobility strategies from lithic reduction patterns (53). In addition, site formation processes observed in the field suggest that the scatters are palimpsests, formed by the gradual incision and erosion of the piedmont surface. It is possible to draw conclusions about patterns of lithic provisioning on the basis of the available information, however.

A systematic archaeological survey in the Karonga district of Malawi, on the other side of the Makutu Range, is instructive (14). The presence of abundant raw material in the form of quartzite cobbles is associated with surface deposits of Early, Middle and Late Stone Age material, dominated by stone tools attributed to the MSA. Thompson and colleagues examined the diversity and distribution of core forms relative to the distribution of cobbles, interpreting the results from a provisioning perspective. Three provisioning strategies were considered: 1) cores are transported as part of a place provisioning strategy, resulting in the accumulation of relatively unexploited cores some distance from raw material sources; 2) cores are transported as part of a personal provisioning strategy, resulting in highly curated cores being deposited near raw material sources; 3) cores are not transported. Thompson and colleagues did not record flakes, although they acknowledge that this could have added to the interpretation of provisioning activities since a fourth possibility is that lithic provisioning was oriented towards the transport of flakes (14). The conclusion reached based on the material from the Karonga survey is that cores were not being transported. This is consistent with excavated material from Chaminade, a dated MSA site in Karonga district, where the percentage of flakes with cortex and of cores with few flake scars is high, which is interpreted as evidence of a flexible strategy of local lithic procurement and proof that people were not just provisioning themselves with raw material at the locality, but were spending time there (51). The abundance of local sources of raw material, therefore, is shown to influence provisioning strategies.

In our sample from northeastern Luangwa, cores occur as isolated artefacts on the landscape and in surface scatters associated with naturally occurring cobble concentrations which functioned as raw material sources. Most of the cores in the surface scatters are not highly curated and we recorded a relatively large number of unmodified flakes (Table 2). The evidence we have gathered to date, therefore, suggests that lithic exploitation strategies in the piedmont zone of the Luwumbu Basin didn’t involve transporting cores (see option 3, above) which we attribute to the frequency with which cobble concentrations formed on the landscape throughout the Stone Age. The occurrence of isolated handaxes and bifacial cores between lithic scatters, however, suggests the provisioning of individuals (option 2). In other words, a flexible strategy of lithic provisioning, involving both the expedient use of local toolstone and individual provisioning, may have prevailed, made possible by the availability of raw materials. A similar situation is described in the Tankwa Karoo, a lowland basin in the interior of South Africa where abundant raw material was exploited expediently during the ESA/MSA, but isolated bifaces and/or handaxes occurring between surface scatters are thought to represent the provisioning of individuals (54).

### Climate controls and chronology

Geomorphological research in the central Luangwa Valley has uncovered evidence of several periods of large-scale instability marked by massive landslides, debris flow and cycles of erosion and redeposition in the eastern piedmont region (8, 9) and in the southern sub-basin (12, 13). This suggests a regional response to large-scale climate shifts, several of which are known to have occurred during the Middle and Upper Pleistocene ((13) citing (55)). Evidence of unconformities and the formation of multiple cobble levels (e.g., at Manzi) suggests that more than one potentially global-scale climate cycle is recorded in the sediment record of the Luangwa Basin. There is a strong possibility, therefore, that the sediment record in the northeastern piedmont region documents a series of major climate events, one of which could be responsible for the formation of the cobble layer at SW23. Whether or not the cycles of flooding and debris flow can be correlated across contexts within the Luangwa valley is unclear, however, and will require better chronological control.

Dating the archaeological material recovered from various locations in the Luangwa Valley directly has proven challenging, e.g., (13). The climate record and its impact on the sedimentary record may offer important clues to the chronology of the deposits that yield archaeological material, however. Climate conditions, mediated by changes in vegetation cover, are the most likely drivers of large-scale landscape transformation in the Luangwa Valley. Climate controls regulate hydrological systems at different scales. Seasonal and inter-annual rainfall patterns regulate the flow of the Luangwa River, periodically altering the course of the numerous seasonally active tributaries flowing into the valley and creating a dynamic floodplain environment (9, 11). Longer-term, alternating cycles of arid and humid conditions, e.g., the African Humid Cycles, also leave their mark in the regional sediment record. Arid cycles result in a thinning of vegetation cover, leaving the landscape more vulnerable to erosion when humid conditions return (55). Since the African Humid cycles are orbitally induced, they provide chronological controls on a millennial scale.

Phytolith analyses (see: Results) indicate that a series of climate fluctuations occurred during the deposition of the sediment package that overlies the cobble layer at SW23 and forms the upper piedmont surface. Sediment cores from Lake Malawi record a shift to wetter conditions in Central Africa during the Middle Pleistocene, followed by a series of aridification cycles (56, 57). Archaeological evidence for human occupation of the piedmont zone during the late ESA and/or early MSA indicates that these climate fluctuations could be related to millennial climate cycles attributable to this timeframe. There have been 216 orbitally induced humid cycles in Africa since the Miocene, however, which means that if we want to tie the events recorded in the sediment sequence in our study region to a specific timeframe, we require better chronological controls.

## Conclusion

The preservation of a sedimentary sequence that overlies the Upper Grit formation in the piedmont zone of northeastern Luangwa highlights the archaeological potential of the study region, which preserves a record of hominin presence spanning the Early to Late Stone Age. A concerted effort to obtain direct dates will be required to establish chronostratigraphic control of the sediment record of northeast Luangwa Valley, without which it will be difficult to reconstruct the history of hominin occupation in the region. It is nevertheless possible to draw some interesting conclusions from the survey results.

The complex history of piedmont formation and subsequent erosion in northeastern Luangwa (see: Discussion) is a confounding factor, resulting in typologically mixed lithic scatters exposed in surface deposits. Despite this, the assemblages described above have a strong MSA signal. The MSA in Africa spans a relatively long timeframe, from ca. 315 ka to the end of the Pleistocene (58, 59). The presence of LCTs, including picks with trihedral points, could also indicate an early post-Acheulean component, possibly dated to the Middle Pleistocene (60). The age of the sediment package from which Stone Age artefacts are eroding in the upper piedmont zone (above 800 meters asl) is currently unknown but, based on the archaeological evidence, it probably post-dates the Miocene when a pattern of global, millennial-scale aridification cycles was established. At SW23, evidence from the phytolith samples is compatible with this suggestion.

Northeastern Luangwa would have been an ecologically attractive environment during the Pleistocene. Topographic controls on regional rainfall patterns governed by orbital forcing will have ensured, then as now, that the valley was relatively well-watered throughout the Pleistocene. Phytolith analyses from the piedmont zone (see: Results) indicate the persistence of miombo forest or dry woodland. This type of environment, characteristic of the Zambezi drainage, is thought to have been one of the most productive environments available to prehistoric hominins in the region (61). Fluvial environments in the monsoon belt, such as the Luangwa Valley, are seasonally important resources today and quite possibly acted as “refugia” during millennial-scale arid climate cycles in the past (62).

The abundance of good-quality lithic raw material in the form of quartz and quartzite cobbles, deposited as fanglomerates during the formation of the piedmont zone and re-exposed by incision, could also explain the persistence of hominin occupation in the region. Patterning in the lithic assemblages suggests a time-transgressive provisioning strategy that includes the expedient exploitation of raw materials at their source and the provisioning of individuals. Finally, in addition to the availability of key resources and an attractive environment, the Luangwa Valley is a natural corridor for human mobility, linking the Central African plateau to the Rift Valley (62). The archaeological potential of the Luangwa Valley would appear to be high, particularly in the piedmont zone of northeastern Luangwa where our fieldwork documents the existence of a thick package of sediments from which Stone Age artefacts are currently eroding. Further fieldwork will allow us to establish the age of this sediment package and confirm that it retains an intact record of hominin occupation.

## Acknowledgements

We would like to thank our Zambian team members for their invaluable assistance with the logistical planning and active participation in the fieldwork. Joseph Museba (National Heritage Conservation Commission, Lusaka) accompanied the team in the field in 2016 and 2019. Martha Nchimunya Kayuni (University of Zambia, Department of Historical and Archaeological Studies) and Margaret Katongo (Assistant Keeper of Archaeology, Livingstone Museum) participated in fieldwork in 2019. Mike Thomas (Professor, Stirling University, retired), Brandon Peecook (Asst. Curator and Professor, Idaho Museum of Natural History & Biological Sciences) and Neil Tabor (Prof. of Earth Sciences, Southern Methodist University), shared their field observations and insights into the geomorphological history of the Luangwa Valley. We also acknowledge the National Heritage Conservation Commission for granting the research permits, and the Livingstone Museum for their assistance and valued support.

## Supporting Information

**SI_1 Phase 1 survey localities.** Topographic and geological attributes of the survey localities.

**SI_2 Sediment analysis : micromorphology.**

**SI_3 Sediment analysis : geochemical grain size analyses**

**SI_4 Phytolith analysis**.

## Reference list

1. Daly MC, Green P, Watts AB, Davies O, Chibesakunda F, Walker R. Tectonics and Landscape of the Central African Plateau and their Implications for a Propagating Southwestern Rift in Africa. Geochemistry, Geophysics, Geosystems. 2020;21(6):e2019GC008746.

2. Peecook BR, Steyer JS, Tabor NJ, Smith RMH. Updated geology and vertebrate paleontology of the Triassic Ntawere Formation of northeastern Zambia, with special emphasis on the archosauromorphs. Journal of Vertebrate Paleontology. 2017;37(sup1):8–38.

3. Sidor CA, Nesbitt SJ. Introduction to vertebrate and climatic evolution in the Triassic Rift Basins of Tanzania and Zambia. Journal of Vertebrate Paleontology. 2017;37(sup1):1–7.

4. MacCrae F, Lancaster DG. 74. Stone Age Sites in Northern Rhodesia. Man. 1937;37:62–4.

5. Clark JD. The Newly Discovered Nachikufu Culture of Northern Rhodesia and the Possible Origin of Certain Elements of the South African Smithfield Culture: Presidential Address. The South African Archaeological Bulletin. 1950;5(19):86–98.

6. Barham L, Phillips WM, Maher BA, Karloukovski V, Duller GAT, Jain M, et al. The dating and interpretation of a Mode 1 site in the Luangwa Valley, Zambia. Journal of Human Evolution. 2011;60(5):549–70.

7. Musonda FB. 100 years of archaeological research in Zambia: changing historical perspectives. South African Archaeological Bulletin. 2012;67(195):88–100.

8. Thomas MF. Evidence for high energy landforming events on the central African plateau: Eastern Province, Zambia. Zeitschrift für Geomorphologie. 1999:273–97.

9. Thomas MF. Geosites and geosystems in quaternary and dynamic landscapes–case studies from a cratonic setting and their wider significance. William Morris Davis–Revista de Geomorfologia. 2021;2(1):1–13.

10. Banks N, Bardwell K, Musiwa S. Karoo rift basins of the Luangwa Valley, Zambia. Geological Society, London, Special Publications. 1995;80(1):285–95.

11. Gilvear D, Winterbottom S, Sichingabula H. Character of channel planform change and meander development: Luangwa River, Zambia. Earth Surface Processes and Landforms. 2000;25(4):421–36.

12. Colton D. An archaeological and geomorphological survey of the Luangwa Valley, Zambia. : BAR publishing; 2009. 210 p.

13. Colton D, Whitfield E, Plater AJ, Duller GAT, Jain M, Barham L. New geomorphological and archaeological evidence for drainage evolution in the Luangwa Valley (Zambia) during the Late Pleistocene. Geomorphology. 2021;392:107923.

14. Thompson JC, Mackay A, de Moor V, Gomani-Chindebvu E. Catchment Survey in the Karonga District: a Landscape-Scale Analysis of Provisioning and Core Reduction Strategies During the Middle Stone Age of Northern Malawi. African Archaeological Review. 2014;31(3):447–78.

15. Roundtree W-ZaM. Integrated Flow Assessment for the Luangwa River. Phase 1: Basin Configuration of EFlows. Lusaka, Zambia; 2018.

16. HydroATLAS-Zambia. Technical Documentation Version 1.0 [Internet]. 2020. Available from: https://www.hydrosheds.org/hydroatlas-zambia.

17. Dixey F. The geomorphology of northern Rhodesia. South African Journal of Geology. 1944;47(01):9–45.

18. Drysdall AR, Kitching JW. A re-examination of the Karoo succession and fossil localities of part of the Upper Luangwa Valley; Réexamen de la succession et des gisements fossilifères du Karroo de la haute vallée de la Luangwa. 1963. p. 62.

19. Peecook BR, Sidor CA, Nesbitt SJ, Smith RM, Steyer JS, Angielczyk KD. A new silesaurid from the upper Ntawere Formation of Zambia (Middle Triassic) demonstrates the rapid diversification of Silesauridae (Avemetatarsalia, Dinosauriformes). Journal of Vertebrate Paleontology. 2013;33(5):1127–37.

20. Sidor CA, Vilhena DA, Angielczyk KD, Huttenlocker AK, Nesbitt SJ, Peecook BR, et al. Provincialization of terrestrial faunas following the end-Permian mass extinction. Proceedings of the National Academy of Sciences. 2013;110(20):8129–33.

21. Colton D. An archaeological and geomorphological survey of the Luangwa Valley, Zambia: University of Liverpool; 2008.

22. Phillipson DW. The Prehistoric Succession in Eastern Zambia: A Preliminary Report. Azania: Archaeological Research in Africa. 1973;8(1):3–24.

23. Elton S, Barham L, Andrews P, Sambrook Smith G. Pliocene femur of Theropithecus from the Luangwa Valley, Zambia. Journal of human evolution. 2003;44(1):133–40.

24. Bishop LC, Barham L, Ditchfield PW, Elton S, Harcourt-Smith WEH, Dawkins P. Quaternary fossil fauna from the Luangwa Valley, Zambia. Journal of Quaternary Science. 2016;31(3):178–90.

25. Chidumayo E. Using natural fertilizers in Miombo woodlands. Issues in African Biodiversity. 1999;2:1–7.

26. Nyirongo VKW. Changes in landuse patterns in upland watersheds of Eastern Luangwa Valley, Zambia, and the potential impact on runoff and erosion. [M.Sc.]: Virginia Polytechnic Institute and State University; 2009.

27. Seifert AV, Vrána S, Kříbek B. Geology of the Muyombe and Luwumbu River areas: explanation of degree sheets 1033 SW and 1033 SE. . Lusaka: Geological Survey Department; 2001.

28. O’Connor E. Report on a visit to Zambia 27 September-4 November 1999. 1999.

29. Vrána S, Kachlík V, Kröner A, Marheine D, Seifert AV, Žáček V, et al. Ubendian basement and its late Mesoproterozoic and early Neoproterozoic structural and metamorphic overprint in northeastern Zambia. Journal of African Earth Sciences. 2004;38(1):1–21.

30. Drysdall AR, Kitching JW. A re-examination of the Karroo succession and fossil localities of part of the Upper Luangwa Valley. Northern Rhodesia, Geological Survey, Memoir. 1963(1).

31. Tabor NJ, Sidor CA, Smith RMH, Nesbitt SJ, Angielczyk KD. Paleosols of the Permian-Triassic: proxies for rainfall, climate change and major changes in terrestrial tetrapod diversity. Journal of Vertebrate Paleontology. 2017;37(sup1):240–53.

32. Angielczyk KD, Steyer J-S, Sidor CA, Smith RM, Whatley RL, Tolan S. Permian and Triassic dicynodont (Therapsida: Anomodontia) faunas of the Luangwa Basin, Zambia: taxonomic update and implications for dicynodont biogeography and biostratigraphy. Early evolutionary history of the Synapsida: Springer; 2014. p. 93–138.

33. Team RC. R: A language and environment for statistical computing. 3.6.3 ed. Vienna, Austria: The R Foundation for Statistical Computing; 2020.

34. Barham LS. The Middle Stone Age of Zambia, South Central Africa: Western academic & specialist Press; 2000.

35. Clark JD, Cormack J, Chin S. Kalambo Falls Prehistoric Site: Volume 3, The Earlier Cultures: Middle and Earlier Stone Age: Cambridge University Press; 1969.

36. Faegri K, Iversen J. Textbook of pollen analysis. 4th ed. Chichester: John Wiley & Sons; 1989.

37. Aleman JC, Saint-Jean A, Leys B, Carcaillet C, Favier C, Bremond L. Estimating phytolith influx in lake sediments. Quaternary Research. 2013;80(2):341–7.

38. Madella M, Alexandre A, Ball T. International code for phytolith nomenclature 1.0. Annals of botany. 2005;96(2):253–60.

39. Scott ICfPTNKkneu-fdSCABTARMVLCL. International code for phytolith nomenclature (ICPN) 2.0. Annals of Botany. 2019;124(2):189–99.

40. Foley R. Off-Site Archaeology: An Alternative Approach for the Short-sited. In: G. Isaac I. Hodder aNH, editor. Pattern of the Past. Cambridge Cambridge University Press; 1981. p. 157–83.

41. Christian A. Tryon. “Early” Middle Stone Age Lithic Technology of the Kapthurin Formation (Kenya). Current Anthropology. 2006;47(2):367–75.

42. Cornelissen E. Site GnJh-17 and its implications for the archaeology of the middle Kapthurin Formation, Baringo, Kenya: Koninklijk Museum voor Midden Afrika; 1992.

43. McBrearty S, Tryon C. From Acheulean to Middle Stone Age in the Kapthurin Formation, Kenya. In: Hovers E, Kuhn SL, editors. Transitions Before the Transition: Evolution and Stability in the Middle Paleolithic and Middle Stone Age. Boston, MA: Springer US; 2006. p. 257–77.

44. Wright DK, Thompson J, Mackay A, Welling M, Forman SL, Price G, et al. Renewed Geoarchaeological Investigations of Mwanganda’s Village (Elephant Butchery Site), Karonga, Malawi. Geoarchaeology. 2014;29(2):98–120.

45. Johnson DL. Biomantle evolution and the redistribution of earth materials and artefacts. Soil Science. 1990;149(2):84–102.

46. Johnson DL, Domier J, Johnson D. Reflections on the nature of soil and its biomantle. Annals of the Association of American Geographers. 2005;95(1):11–31.

47. Retallack GJ. Soils of the past: an introduction to paleopedology: John Wiley & Sons; 2008.

48. Rothwell RG, Croudace Iw. Twenty Years of XRF Core Scanning Marine Sediments: What Do Geochemical Proxies Tell Us? In: Croudace IW, Rothwell RG, editors. Micro-XRF Studies of Sediment Cores: Applications of a non-destructive tool for the environmental sciences. Dordrecht: Springer Netherlands; 2015. p. 25–102.

49. Van Hoang L, Clift PD, Schwab AM, Huuse M, Nguyen DA, Zhen S. Large-scale erosional response of SE Asia to monsoon evolution reconstructed from sedimentary records of the Song Hong-Yinggehai and Qiongdongnan basins, South China Sea. Geological Society, London, Special Publications. 2010;342(1):219–44.

50. Fanning PC, Holdaway SJ, Rhodes EJ, Bryant TG. The surface archaeological record in arid Australia: Geomorphic controls on preservation, exposure, and visibility. Geoarchaeology. 2009;24(2):121–46.

51. Nightingale S, Schilt F, Thompson JC, Wright DK, Forman S, Mercader J, et al. Late Middle Stone Age Behavior and Environments at Chaminade I (Karonga, Malawi). Journal of Paleolithic Archaeology. 2019;2(3):258–97.

52. Bicho N, Haws J, Raja M, Madime O, Gonçalves C, Cascalheira J, et al. Middle and Late Stone Age of the Niassa region, northern Mozambique. Preliminary results. Quaternary International. 2016;404:87–99.

53. Riel-Salvatore J, Barton CM. Late Pleistocene technology, economic behavior, and land-use dynamics in southern Italy. American Antiquity. 2004:257–74.

54. Hallinan E. Landscape-scale perspectives on Stone Age behavioural change from the Tankwa Karoo, South Africa. Azania: Archaeological Research in Africa. 2021;56(3):304–43.

55. Thomas MF. Landscape sensitivity to rapid environmental change—a Quaternary perspective with examples from tropical areas. CATENA. 2004;55(2):107–24.

56. Scholz CA, Cohen AS, Johnson TC, King J, Talbot MR, Brown ET. Scientific drilling in the Great Rift Valley: The 2005 Lake Malawi Scientific Drilling Project — An overview of the past 145,000years of climate variability in Southern Hemisphere East Africa. Palaeogeography, Palaeoclimatology, Palaeoecology. 2011;303(1):3–19.

57. Lyons RP, Scholz CA, Cohen AS, King JW, Brown ET, Ivory SJ, et al. Continuous 1.3- million-year record of East African hydroclimate, and implications for patterns of evolution and biodiversity. Proceedings of the National Academy of Sciences. 2015;112(51):15568–73.

58. Richter D, Grün R, Joannes-Boyau R, Steele TE, Amani F, Rué M, et al. The age of the hominin fossils from Jebel Irhoud, Morocco, and the origins of the Middle Stone Age. Nature. 2017;546(7657):293–6.

59. Scerri EML. The North African Middle Stone Age and its place in recent human evolution. Evolutionary Anthropology: Issues, News, and Reviews. 2017;26(3):119–35.

60. Brooks AS, Yellen JE, Potts R, Behrensmeyer AK, Deino AL, Leslie DE, et al. Long-distance stone transport and pigment use in the earliest Middle Stone Age. Science. 2018;360(6384):90–4.

61. Barham LS, Brown KR. Central Africa and the emergence of regional identity in the Middle Pleistocene. Human Roots: Africa and Asia in the Middle Pleistocene: Western Academic and Specialist Press; 2001. p. 65–80.

62. Burrough SL, Thomas DSG, Barham LS. Implications of a new chronology for the interpretation of the Middle and Later Stone Age of the upper Zambezi Valley. Journal of Archaeological Science: Reports. 2019;23:376–89.

